## Supplementary Figures and Tables for "Applying 3D correlative structured illumination microscopy and X-ray tomography to characterise herpes simplex virus-1 morphogenesis"

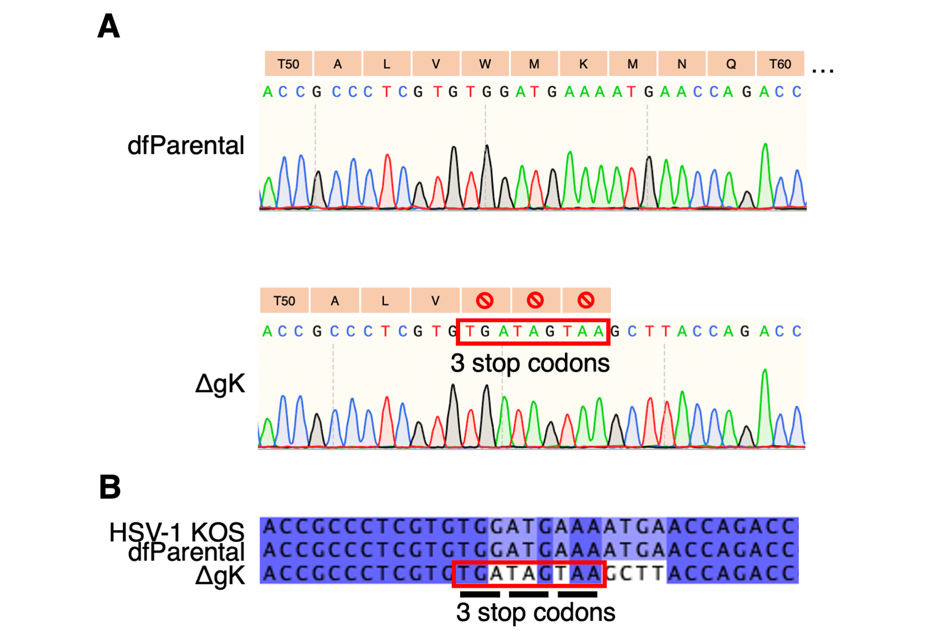

**Supplementary Figure 1. Sequence alignment of UL53 gene in ΔgK and dfParental HSV-1.** (**A**) Sanger sequencing confirmed the presence of the three in-frame stop codons in the UL53 gene (encoding gK) of the ΔgK virus. Translated amino acid sequences are shown. (**B**) UL53 sequences from ΔgK and dfParental HSV-1 were aligned with the UL53 sequence of the KOS genome (GenBank accession number NC_001806)^104^.

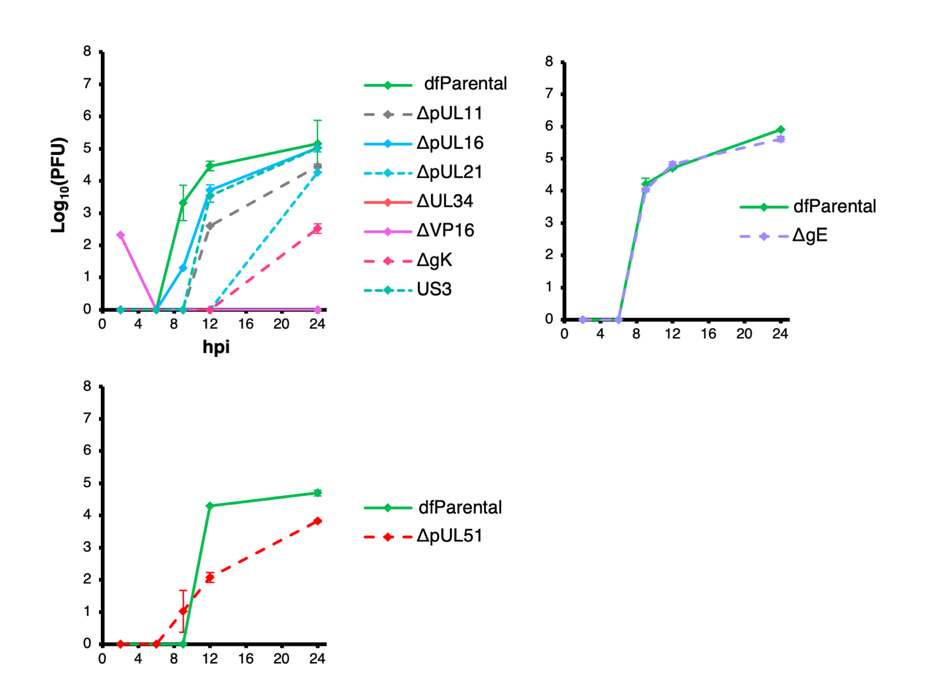

**Supplementary Figure 2. Additional replicate for single-step replication kinetics of mutant and dfParental HSV-1**. Single-step replication curves on U2OS cells infected at MOI = 2 with virus. dfParental refers to the parental eYFP-VP26 & gM-mCherry KOS strain. U2OS cells were infected at MOI = 2 with virus over a 24 h period and were treated with citric acid at the 1-hour timepoint to deactivate extracellular viruses. Titrations were performed on parental or complementing Vero cells. Two technical repeats were measured for each timepoint, and the data are representative of two biological replicates (**Fig. 2**). Error bars show mean ± range.

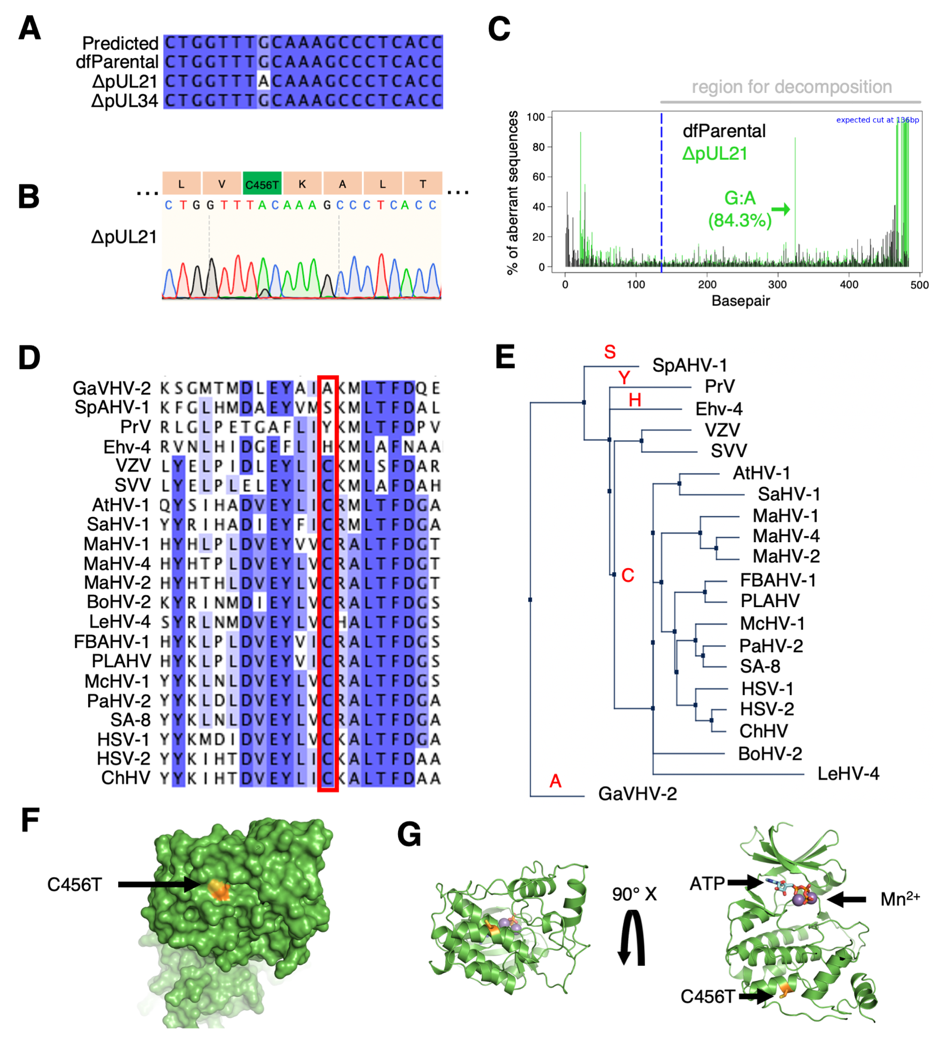

**Supplementary Figure 3. Sequence alignment of US3 in recombinant and dfParental viruses.** (**A**) Sanger sequencing of the US3 gene revealed a guanine to adenine mutation at base pair 1,367 in the ΔpUL21 virus but not the ΔUL34 virus. Sequencing results were compared against the KOS genome (HSV-1 KOS; GenBank accession number NC_001806)^104^. (**B**) This mutation resulted in a C456T substitution in pUS3. (**C**) Tracking of Indels by Decomposition (TIDE)^107^ analysis was performed to identify prevalent base pair changes in the US3 gene of the ΔpUL21 virus. The mutation at base pair 1,367 was 84.3% prevalent and no other prevalent mutations were identified. (**D**) pUS3 orthologues from 21 alphaherpesviruses were aligned to assess conservation of amino acid residue in question. Only four species contained different residues at this position. Species aligned (abbreviation, sequence ID): Gallus alphaherpesvirus 2 (GaVHV-2, GenBank: ACF94893.1), Spheniscid alphaherpesvirus 1 (SpAHV-1, GenBank: SCL76985.1), Pseudorabies virus (PrV, UniProtKB/Swiss-Prot: P24381.2), Equine herpesvirus 4 (Ehv-4, GenBank: BAV93592.1), Varicella Zoster Virus (VZV, NCBI Reference Sequence: NP_040188.1), Simian Varicella Virus (SVV, UniProtKB/Swiss-Prot: Q04543.1), Ateline herpesvirus 1 (AtHV-1, NCBI Reference Sequence: YP_009361942.1), Saimiriine herpesvirus 1 (SaHV-1, NCBI Reference Sequence: YP_003933845.1), Macropodid alphaherpesvirus 1 (MaHV-1, NCBI Reference Sequence: YP_009227221.1), Macropodid alphaherpesvirus 4 (MaHV-4, NCBI Reference Sequence: YP_010801716.1), Macropodid alphaherpesvirus 2 (MaHV-2, NCBI Reference Sequence: YP_010798801.1), Bovine alphaherpesvirus 2 (BoHV-2, NCBI Reference Sequence: YP_010798781.1), Leporid alphaherpesvirus 4 (LeHV-4, NCBI Reference Sequence: YP_009230196.1), Fruit bat alphaherpesvirus-1 (FBAHV-1, NCBI Reference Sequence: YP_009042124.1), Pteropus lylei-associated alphaherpesvirus (PLAHV, NCBI Reference Sequence: YP_010801544.1), Macacine alphaherpesvirus 1 (McHV-1, NCBI Reference Sequence: YP_010797361.1), Papiine alphaherpesvirus 2 (PaHV-2, GenBank: AHM96184.1), Herpes simian agent 8 (SA-8, NCBI Reference Sequence: YP_164505.1), Herpes simplex virus 1 (HSV-1, GenBank: AKG59533.1), Herpes simplex virus 2 (HSV-2, UniProtKB/Swiss-Prot: P13287.1), Chimpanzee herpesvirus (ChHV, NCBI Reference Sequence: YP_009011050.1) (**E**) A neighbour-joining tree of the aligned sequences revealed the pUS3 orthologues containing the conserved C residue were more similar to each other than to the orthologues containing the A, H, S, or Y residues. (**F**) A model of pUS3 was superposed onto the structure of Protein Kinase A^33^, and a surface map revealed the mutated residue is exposed. (**G**) A ribbon model of the structure in F shows that the mutated residue is not located near the kinase active site, which is indicated by the ATP and metal ions^123^.

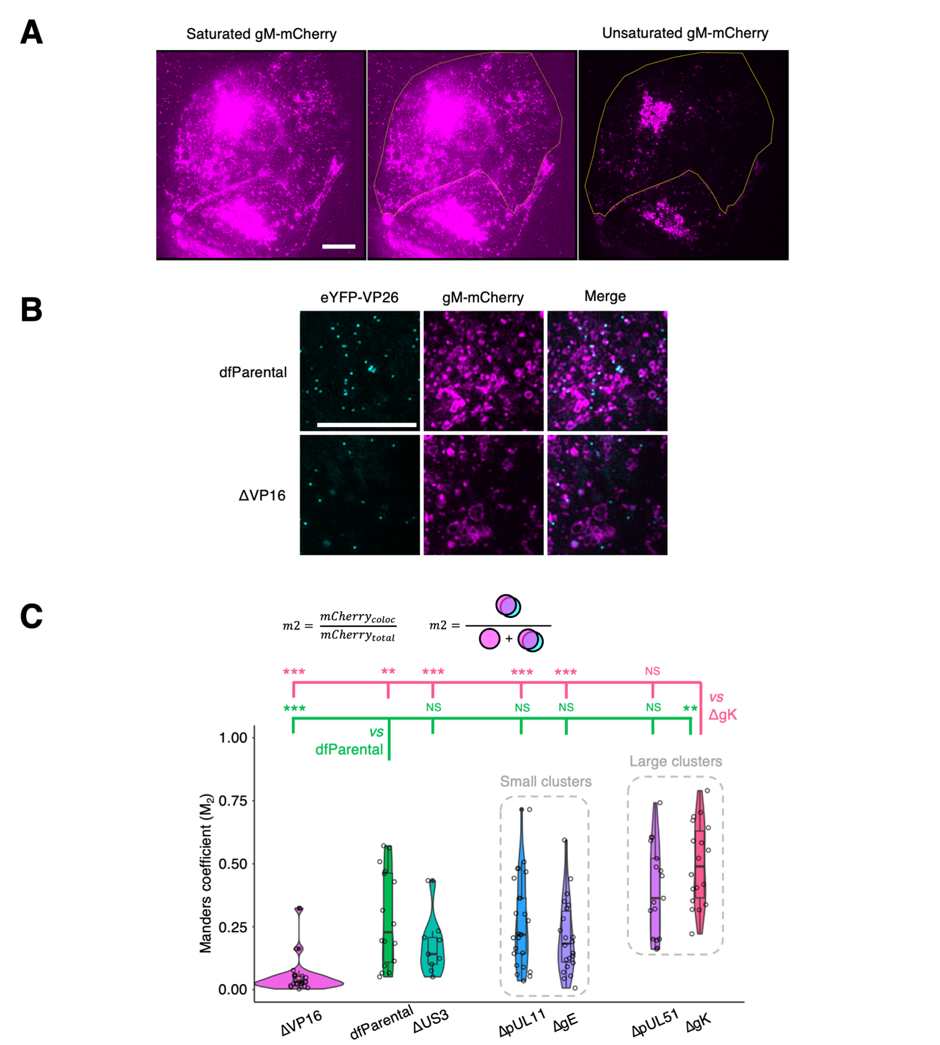

**Supplementary Figure 4. Association of gM-mCherry+ endomembranes with capsids.** U2OS cells were infected at MOI = 2 with indicated viruses for 16 hours. Scale bars = 10 μm. (**A**) Saturation of gM-mCherry^+^ endomembranes was used to delineate the borders of the cytoplasm (yellow silhouette). This example taken from Fig. 4A shows a cell infected with the dfParental virus. (**B**) Individual and combined channels for data in Fig. 5B showcasing the reduced association of ΔVP16 capsids with gM-Cherry^+^ endomembranes compared with the dfParental virus. (**C**) Manders coefficients (M_2_) were measured for each virus and represent the amount of gM-mCherry fluorescence that colocalised with eYFP-VP26 (mCherry_coloc_) as a proportion of total gM-mCherry fluorescence at the JAC (mCherry_total_). M_2_ values were markedly lower for the ΔVP16 virus than other viruses. Data from the ΔUS3 virus were included as negative control for attenuation in cytoplasmic virion assembly. Mann-Whitney *U* tests were performed to assess significance of differences between dfParental (N=16) (green statistics) or ΔgK (N=18) (pink statistics) and other viruses, specifically ΔUS3 (N=9), ΔVP16 (N=22), ΔgE (N=23), ΔpUL11 (N=25), and ΔpUL51 (N=17). P-value thresholds: <0.05 (*), <0.005 (**), and <0.0005 (***). NS, no significance.

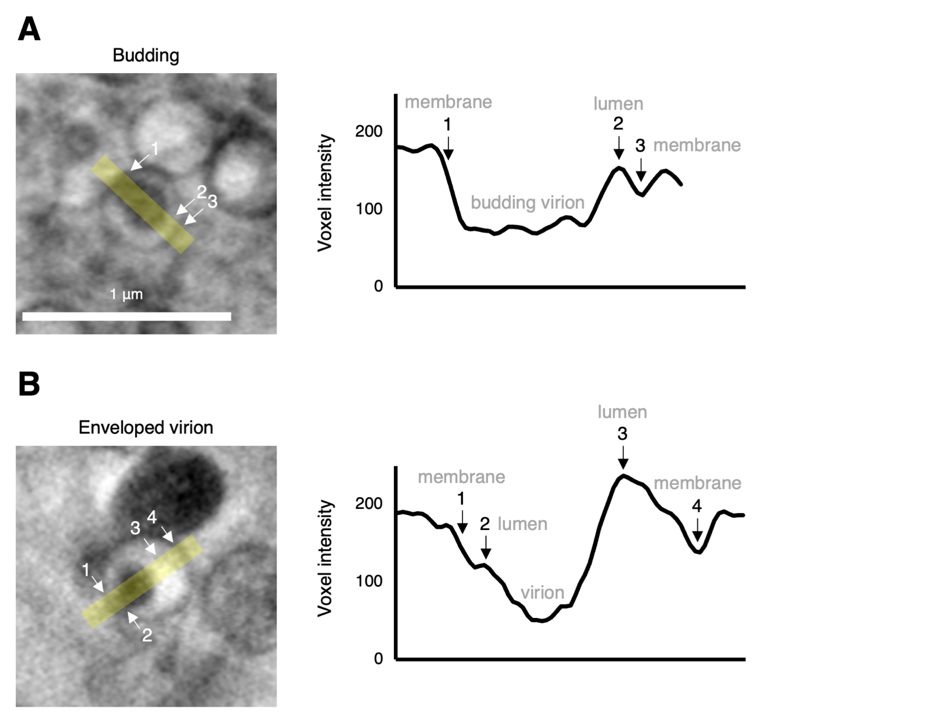

**Supplementary Figure 5. Full and partial envelopment intermediates distinguished based on voxel intensity.** Using U2OS cells infected at MOI = 2 with the ΔpUL51 virus for 16 hours, cryoSXT captures (**A**) a virus particle budding into the lumen of a vesicle and (**B**) a fully enveloped virion within a vesicle. Voxel intensities were measured from the yellow stripes (width = 10 voxels) using Fiji, and notable features are numbered. The budding virion in A cannot be distinguished from the membrane, whereas the virion in B is separated from the membrane by a brighter lumen. Scale bar = 1 μm.

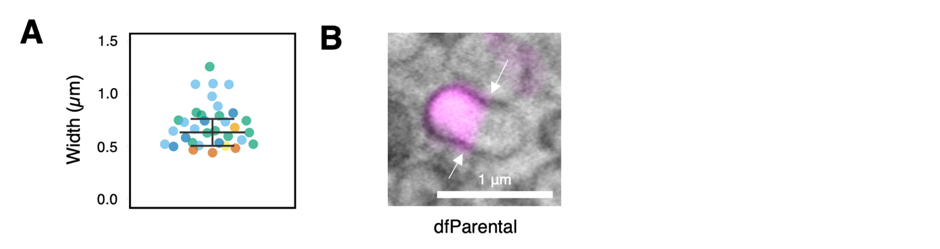

**Supplementary Figure 6. Features of gM-mCherry^+^ vesicles in infected cells.** (**A**) Widths of vesicles associated with capsid arrays were measured using *Contour*^68^ and are colour-coded by source tomogram. Error bars show mean ± SD (n = 34). (**B**) Polarisation of gM-mCherry was also observed in dfParental-infected cells and membrane constrictions were visible (arrows), suggesting vesicle fission, fusion, or pressure imposed by microtubules. Scale bar = 1 μm.

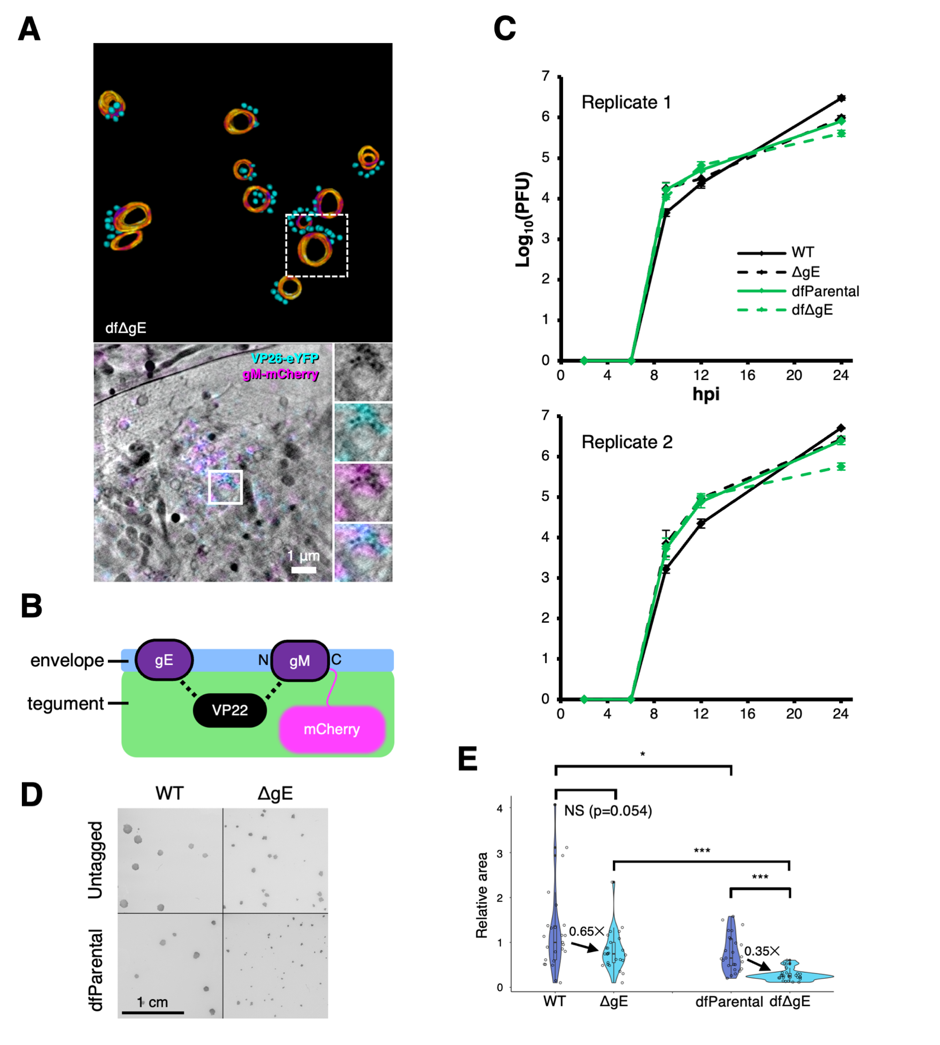

**Supplementary Figure 7. Impact of the gM-mCherry conjugation on the ΔgE virus.** (**A**) Numerous stalled envelopment events were observed for the dual fluorescent ΔgE virus (labelled dfΔgE only in this figure). Scale bar = 1 μm. (**B**) Schematic of known interactions between tegument component VP22 and envelope components gE and gM. (**C**) Neither absence of gE expression nor gM-mCherry tagging dramatically altered the replication kinetics of HSV-1 mutants. U2OS cells were infected at MOI = 2 with virus over a 24 h period and were treated with citric acid at the 1-hour timepoint to deactivate residual input viruses. Two biological replicates were performed, with two technical repeats for each timepoint. dfParental and dfΔgE data are the same as those shown in **Fig. 2A** and **Supp. Fig. 2**. Error bars show mean ± SD. (**D**) U2OS cells were infected with indicated viruses, were covered in media containing 0.6% (v/v) CMC after 1 hour, and plaques were fixed and stained 72 hpi. Scale bar = 1 cm. (**E**) Relative plaque size was calculated for each virus. Mann-Whitney *U* tests were used to assess the significance of differences. P-value thresholds: <0.05 (*), <0.005 (**), and <0.0005 (***). NS, no significance.

**Supplementary Table 1. Primers to generate mutants by two-step red recombination.** Sequences homologous to the gene are in upper case, sequences homologous to the template plasmids pEP-KanS or pEP-RenillaLuc are in lower case, mutated start codons are underlined, and introduced stop codons are in bold.

| **Gene** | **Forward primer (5′ to 3′)** | **Reverse primer (5′ to 3′)** |
| --- | --- | --- |
| UL11 | GAAGCAGCCGCCTGGCGTTCGACGACACGCTCGCCGAGCTctgggc**tag**TCGTTCTCCGGGACCCGGCCaggatgacgacgataagtaggg | CGTTGTTTCGGCAGCAGCAGGGCCGGGTCCCGGAGAACGA**cta**gcccagAGCTCGGCGAGCGTGTCGTCcaaccaattaaccaattctgattag |
| UL16 | GGCGCAGCTGGGACCCCGGCGGCCCCTGGCGCCGCCTGGT**TGATAGTAA**GCTTGCCCCGGCCGGATTCCCaggatgacgacgataagtaggg | CGTGCCGCGAGCTCCGGCCCGGGAATCCGGCCGGGGCAAGC**TTACTATCA**ACCAGGCGGCGCCAGGGGCCcaaccaattaaccaattctgattag |
| UL21 | GCACTACCGGGACGTTGTGTTTTACGTCACAACGGACCGA**TGATAGTAA**GCTTTGTGTGCGGGGGGTGTGaggatgacgacgataagtaggg | CGGCCGCCCCACGGAATAAACACACCCCCCGCACACAAAGC**TTACTATCA**TCGGTCCGTTGTGACGTAAAcaaccaattaaccaattctgattag |
| UL34 | CCCTTTGGTGGGTTTACGCGGGCACGCACGCTCCCATCGCGGGCGCCatgacttcgaaagtttatgatcc | GCTTAAGACCCCGCAGGGCCTGGTGCCACGGGCGGGAGGGCCCTTGGGTTTTAttgttcatttttgagaactcgc |
| UL48 | CAAAAGCCCGATATCGTCTTTCCCGTATCAACCCCACCCAGAATTCTTTACCGATGCCCTTGGAATaggatgacgacgataagtaggg | CCTACCCACCGTACTCGTCAATTCCAAGGGCATCGGTAAAGAATTCTGGGTGGGGTTGATACGGGAcaaccaattaaccaattctgattag |
| UL51 | TATATGTGGCTGGGGAGCGCGCCCCGAGGAACAATATGAG**TAGTGATAA**GGATCCGTTCCGCCCTCGGAGGCGGAaggatgacgacgataagtaggg | GGGCCTCCTGCAGCCGCGGCTCCGCCTCCGAGGGCGGAACGGATCC**TTATCACTA**CTCATATTGTTCCTCGGGGCcaaccaattaaccaattctgattag |
| UL53 | GGTACGCCCCACCGGCACCAACAACGACACCGCCCTCGTG**TGATAGTAA**GCTTACCAGACCCTATTGTTTCTGaggatgacgacgataagtaggg | GGGGGGTGCGTCGGGGCCCCCAGAAACAATAGGGTCTGGTAAGC**TTACTATCA**CACGAGGGCGGTGTCGTTGTcaaccaattaaccaattctgattag |
| US3 | CACCACACCACCCGGCGATGCCGAGCGCCTGTGTCATCTGTGATCTTCGAGACTGCCGTCaggatgacgacgataagtaggg | GAGAACAAGGACGCGTTGTGGACGGCAGTCTCGAAGATCACAGATGACACAGGCGCTCGGcaaccaattaaccaattctgattag |
| US8 | GGGGTTTCTTCTCGGTGTTTGTGTTGTATCGTGCTTGGCG**TAGTGATAA**GCTTCGTCCTGGAGACGGGTGAGTaggatgacgacgataagtaggg | AACGAAACGTCCTCGCCGACACTCACCCGTCTCCAGGACGAAGC**TTATCACTA**CGCCAAGCACGATACAACACcaaccaattaaccaattctgattag |

**Supplementary Table 2. PCR amplification primers for HSV-1 US3 and UL53.**

| **Gene** | **Forward primer (5′ to 3′)** | **Reverse primer (5′ to 3′)** |
| --- | --- | --- |
| UL53 | GGTCCTCCTACAGCTAGTCC | GCTGGGTTGGTCTTGGTAAC |
| US3 | CAGATTTGTAAGGCCACGCAC | GATAATGGGAACAACGGCACG |

**Supplementary Table 3. Sanger sequencing primers for HSV-1 US3 and UL53.**

| **Gene** | **Complementary strand** | **Primer (5′ to 3′)** |
| --- | --- | --- |
| UL53 | Sense | CAAATGCGACAGCAACCG |
|  | Sense | CTCTTGAACTACGCAGGC |
| US3 | Sense | CTGCCGCTCCTTAAAACC |
|  | Sense | CAGAAGAGCTGGACGCCATG |
|  | Sense | GATCAAGCCCCTTCCCCTAC |
|  | Sense | CCACCGCGACATTAAGAC |
|  | Anti-sense | GTGATCTGACTGTCGCACG |
